# Development of a novel signature of long noncoding RNAs as a prognostic biomarker for esophageal cancer

**DOI:** 10.1101/441568

**Authors:** Yan Miao, Jing Sui, Ying Zhang, Lihong Yin

## Abstract

**Objectives:** This study aims to develop a lncRNA signature based on RNA-Seq data to predict overall survival in esophageal cancer patients.

**Methods:** The lncRNA expression profiles and clinical data were downloaded from The Cancer Genome Atlas (TCGA) database on August 30, 2017. Differentially expressed lncRNAs were screened out between tumor tissues and adjacent normal tissues. The univariate and multivariate Cox regression models were used to develop a prognostic signature for all esophageal cancer patients. The receiver operating curve (ROC) was used to test the sensitivity and specificity of lncRNA signature. Survivals were compared via log-rank test. GO and KEGG enrichment analyses were used to explore the potential functions of prognostic lncRNAs.

**Results:** We identified two lncRNAs (RPL34-AS1 and GK3P) were significantly associated with the overall survival of the total 150 esophageal cancer patients. A novel two-lncRNA signature was constructed by Cox regression models. Signature low-risk cases showed better overall survival (median 625.560 days vs. 478.000 days, *p* = 0.002) than high-risk cases. Further analysis suggested that this two-lncRNA signature was independent of clinical characteristics. GO functional and KEGG pathway enrichment analyses revealed potential functional roles of the two prognostic lncRNAs in tumorigenesis.

**Conclusions:** Our findings suggest that the two-lncRNA signature may be a useful prognostic biomarker for predicting overall survival in esophageal cancer patients.

## INTRODUCTION

Esophageal cancer (EC) ranks the eighth most common cancer worldwide and the sixth leading cause of cancer-related mortality (Ferlay et al. 2015). Unfortunately, EC is often associated with a poor prognosis that having an 18% overall five-year survival (Malhotra et al. 2017). As with most solid tumors, pathological tumor-node-metastasis (TNM) staging is still the primary indicator of survival in patients with EC (Malhotra et al. 2017; Rice et al. 2017). The molecular heterogeneity and complexity of EC make it difficult to predict its clinical outcomes (Hao et al. 2016; Hardie et al. 2005). Therefore, it is urgent to find specific prognostic factors of EC which would improve treatment efficacies for patients.

In the age of genomics, researchers have discovered tens of thousands of long noncoding RNAs (lncRNAs) by genome-wide techniques, which are defined as RNA transcripts exceeding 200 nucleotides in length without protein-coding capacity (Schmitt & Chang 2016; Young & Ponting 2013). Emerging evidence revealed that lncRNAs play a critical role in various aspects of cellular processes, including proliferation, migration, invasion, and genomic stability (Iyer et al. 2015; Ling et al. 2015). Based on the expression specificity among different tumor types, lncRNAs could serve as the basis for many clinical applications in oncology (Li & Chen 2013; Shi et al. 2013; Wahlestedt 2013). Till now, lncRNA biomarkers for diagnosis of EC have been reported in many studies (Tong et al. 2015; Wang et al. 2017; Wu et al. 2015). However, the understanding of the prognostic value of lncRNAs in patients with EC from previous studies was limited.

The Cancer Genome Atlas (TCGA) is a database, which provides a collection of DNA copy-number variation, DNA methylation, RNA sequence, and corresponding clinical data for the top 25 tumor types. In this study, we used the RNA-Seq dataset from the TCGA database to identify candidate prognostic lncRNAs for EC. After integrative analysis, we developed a prognostic lncRNA signature for the overall survival (OS) prediction in EC patients.

## MATERIALS & METHODS

### TCGA data source

185 EC patients’ data were downloaded from the TCGA database on August 30, 2017. The exclusion criteria were listed in the following: i) histologic diagnosis ruled out EC; ii) another malignancy besides EC. After that, 150 EC patients were included in this study. As the data were downloaded from the public database, ethical approval was not applicable here. Data processing procedures followed the guidelines of TCGA data access (http://cancergenome.nih.gov/publications/publicationguidelines).

### LncRNA data mining and processing

The level 3 RNA-Seq data were obtained from the TCGA database. Raw data of lncRNA sequencing were processed and normalized by TCGA RNASeqv2 system (Zand et al. 2016). Here, only lncRNAs with the description in NCBI (http://ncbi.nlm.nih.gov/gene) or Ensembl (http://ensembl.org/index.html) were selected for further analysis. To identify differentially expressed lncRNAs, the expression of lncRNAs in tumor tissues were screened and compared with that in adjacent non-tumor tissues. The candidate lncRNAs were gathered for further analysis.

### Construction of the prognostic signature

The expression profile of each lncRNA was normalized by log2 transformation for statistical analysis. The univariate Cox regression model was used to evaluate the EC-specific lncRNAs that were associated with OS. The multivariate Cox regression model was used to evaluate the prognostic value of these OS-related lncRNAs. The risk-score model was constructed based on the combination of the expression profiles of each prognostic lncRNA, weighted by the regression coefficients that were calculated by multivariate Cox regression analysis. The formula was listed as follows: 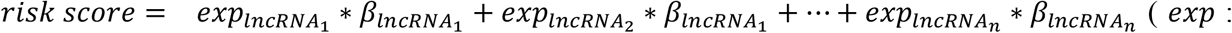 expression level; *β*: regression coefficient derived from the multivariate Cox regression model) (Zeng et al. 2017). The median risk score was set to the cutoff point, and EC patients were divided into high or low groups (Zhou et al. 2016). Further univariate and multivariate Cox regression models were used to investigate the effects of risk score and clinical characteristics in EC patients. The hazard ratio (HR) and the 95% confidence interval (CI) were used for quantification. The time-dependent receiver operating curve (ROC) within five years was used to assess the predictive value of risk score for time-dependent outcomes (Heagerty et al. 2000). The Kaplan-Meier (K-M) curve and log-rank test were used to assess the differences in survival distribution among different groups. Here, the *p* value <0.05 was considered to be significant. IBM SPSS Statistics 22.0 (IBM Corp., Armonk, NY, USA) was used to perform the analyses mentioned above.

### Functional enrichment analysis

To investigate the biological features of these two lncRNAs, we selected the genes that were strongly correlated with these two lncRNAs expressions (Pearson |R| > 0.5) from TCGA database. Pathways and biological processes were predicted by functional enrichment analysis of the Kyoto Encyclopedia of Genes and Genomes (KEGG) and Gene Ontology (GO) in the DAVID database (http://david.ncifcrf.gov/). The *p*-value <0.05 was considered to be significant. Afterward, the protein-protein interaction (PPI) network was constructed based on the co-expressed genes via the STRING database (http://string-db.org/).

## RESULTS

### Patient characteristics

There were 150 EC patients included in this study. According to the cancer staging standards of the American Joint Committee on Cancer (AJCC) (Amin & Edge 2017), the EC patients were classified into four groups: stage I (16 cases), stage II (74 cases), stage III (52 cases), and stage IV (8 cases). The mean (and standard deviation) age of all EC patients was 61.233 (± 11.311) years. The mean OS was 551.780 (± 505.185) days. 61 of 150 (40.667%) patients died during follow-up.

### Identification of differentially expressed lncRNAs

269 lncRNAs were identified as differentially expressed in EC tissues and adjacent non-tumor tissues, including 152 lncRNAs in stage I, 219 lncRNAs in stage II, 183 lncRNAs in stage III, and 109 lncRNAs in stage IV. We used “Fold change > 2 or < 0.5, *p* < 0.05,and FDR < 0.05” as the screening method to identify the significant differentially expressed lncRNAs. Then, 74 candidate lncRNAs were identified as differentially expressed in all tumor stages (Fig. 1 and Fig. 2).

**Figure 1.**
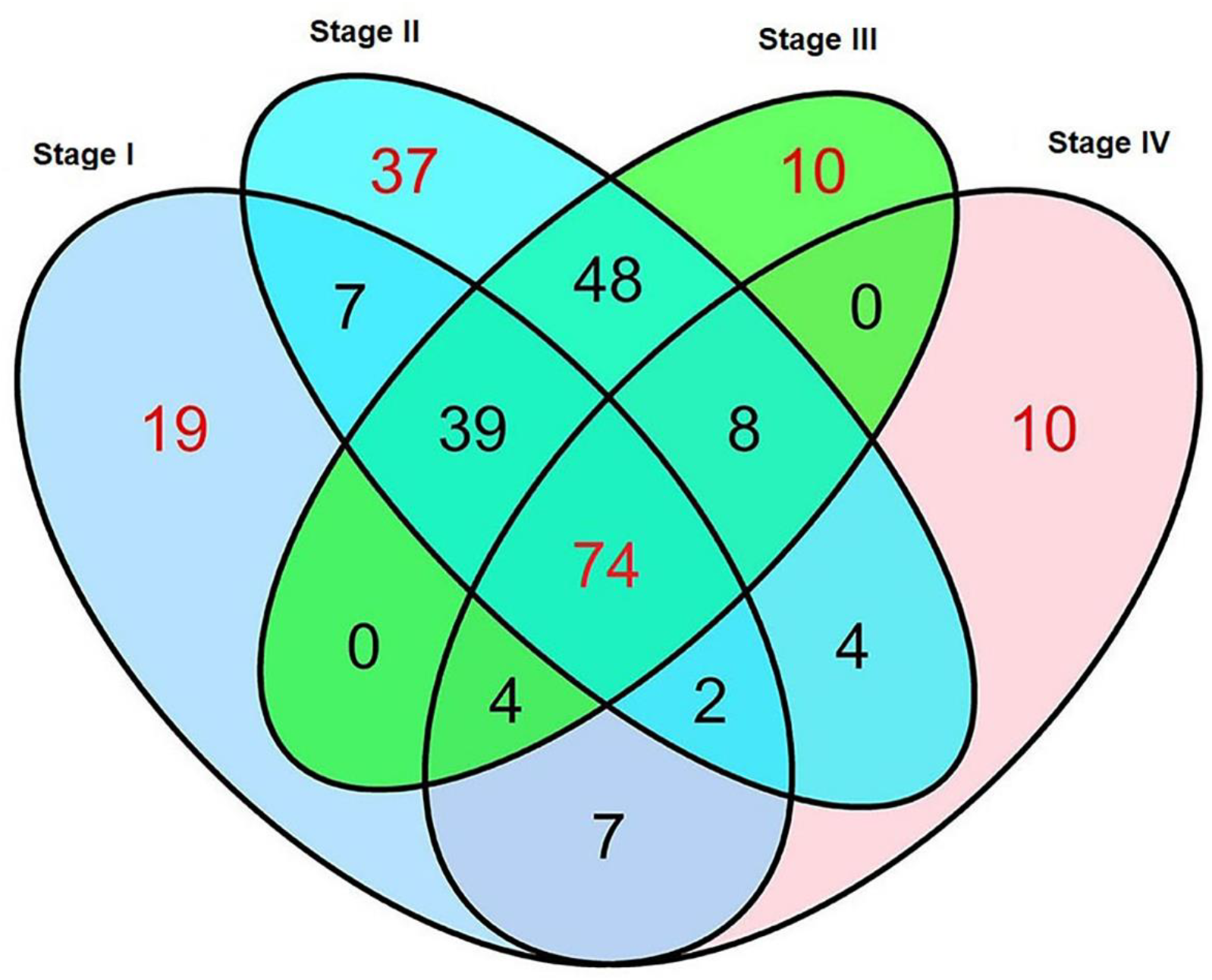
Venn diagram analysis of differentially expressed lncRNA in esophageal cancer.

**Figure 2.**
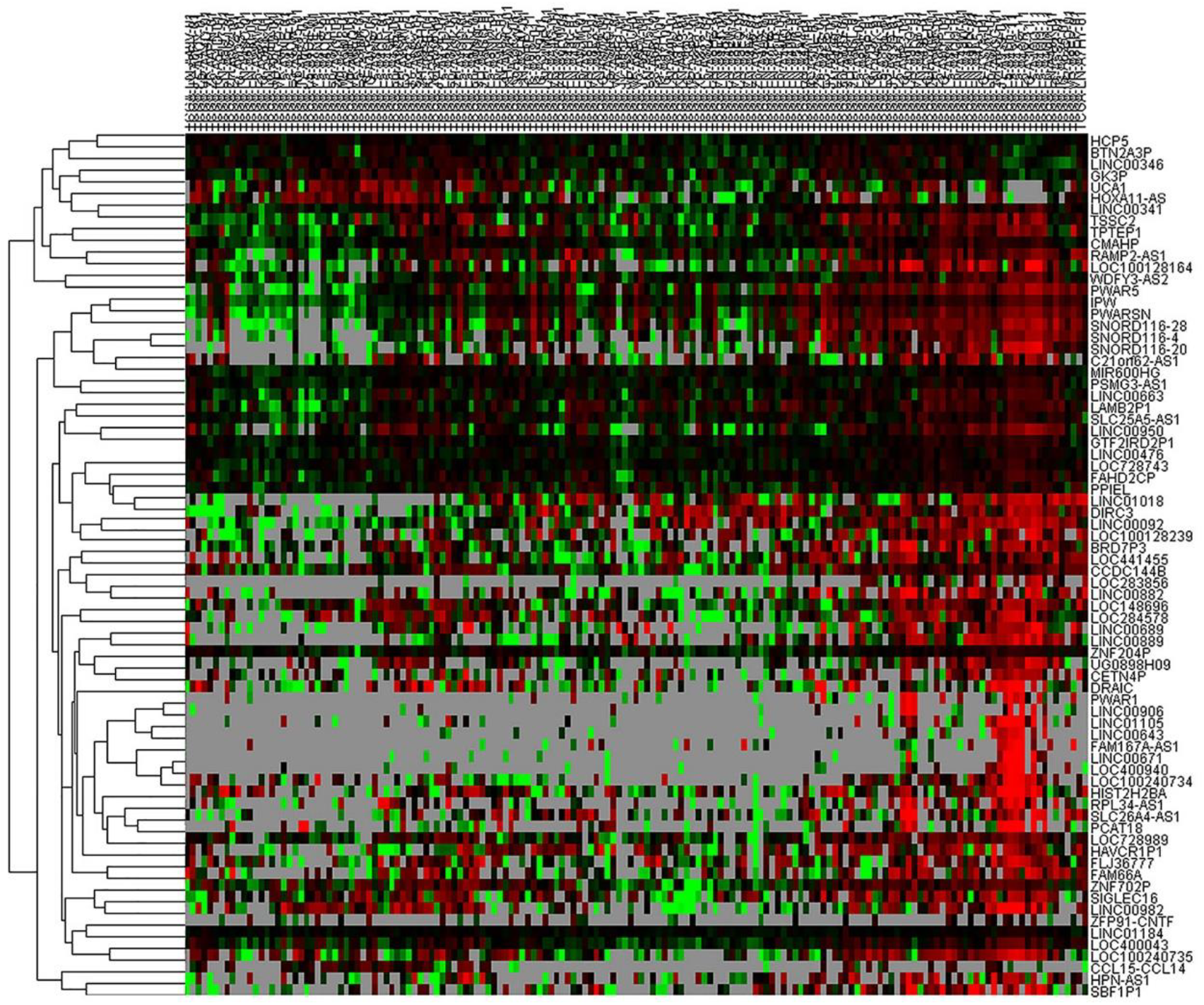
Heatmap of 74 lncRNAs differentially expressed in esophageal cancer.

### Construction of a prognostic signature for EC patients

Based on the 74 lncRNAs and the clinical information extracted from TCGA database, two candidate lncRNAs were identified to be significantly associated with OS (*p* < 0.05) in EC patients by univariate Cox regression analysis (Table 1). The multivariate Cox regression analysis was used to verify the correlation between two candidate lncRNAs and OS (Table 1 and Fig 3A). These two lncRNAs, RPL34-AS1 and GK3P, also showed prognostic value for EC patients (Fig 3B).

**Figure 3.**
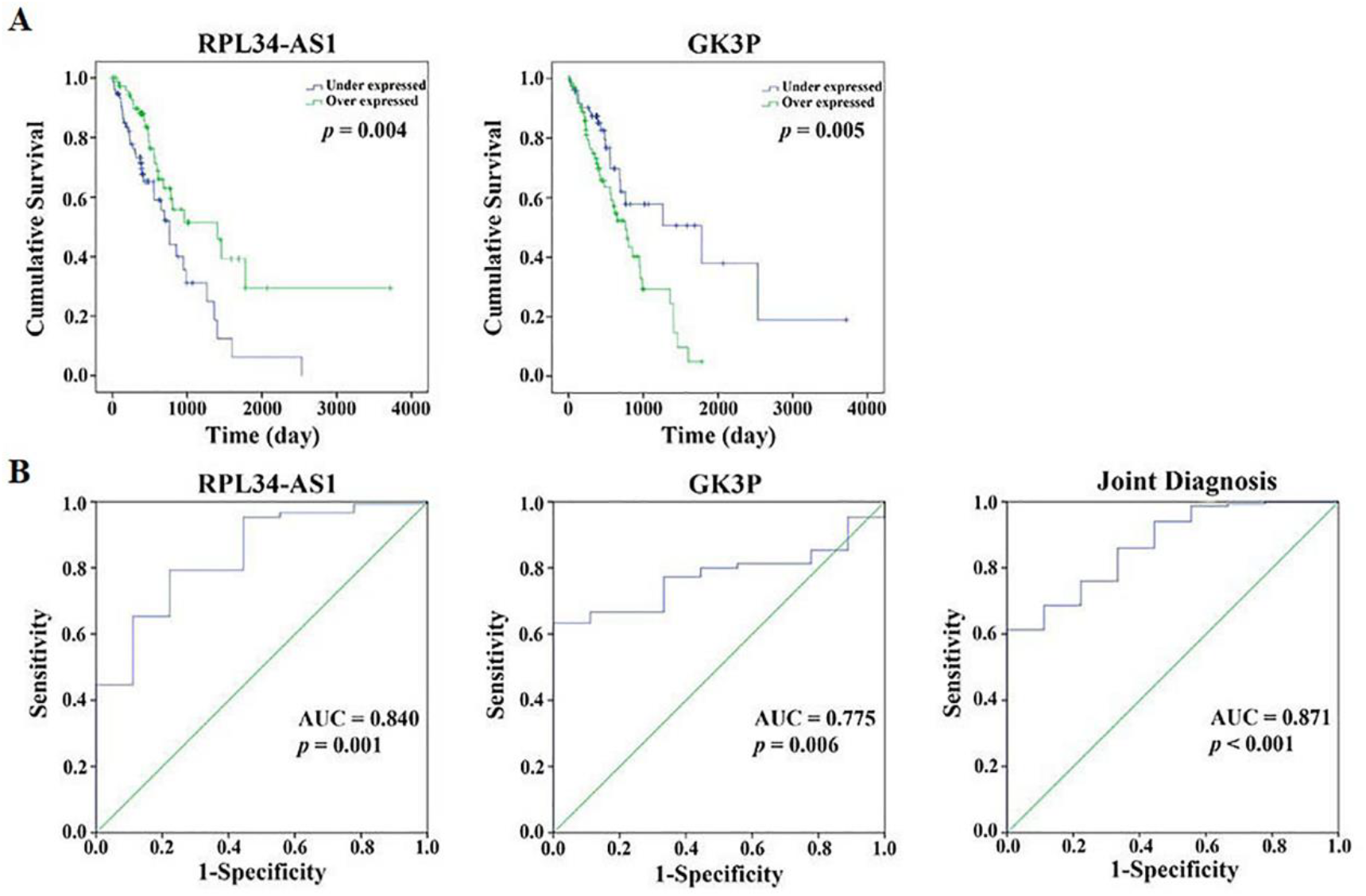
Two differentially expressed lncRNAs in esophageal cancer patients. (A) Kaplan-Meier curves show the correlation between two lncRNAs and overall survival; (B) ROC curves compare the sensitivity, specificity, and prognostic value of two lncRNAs and assess the efficacy of the signature.

**Table 1.**
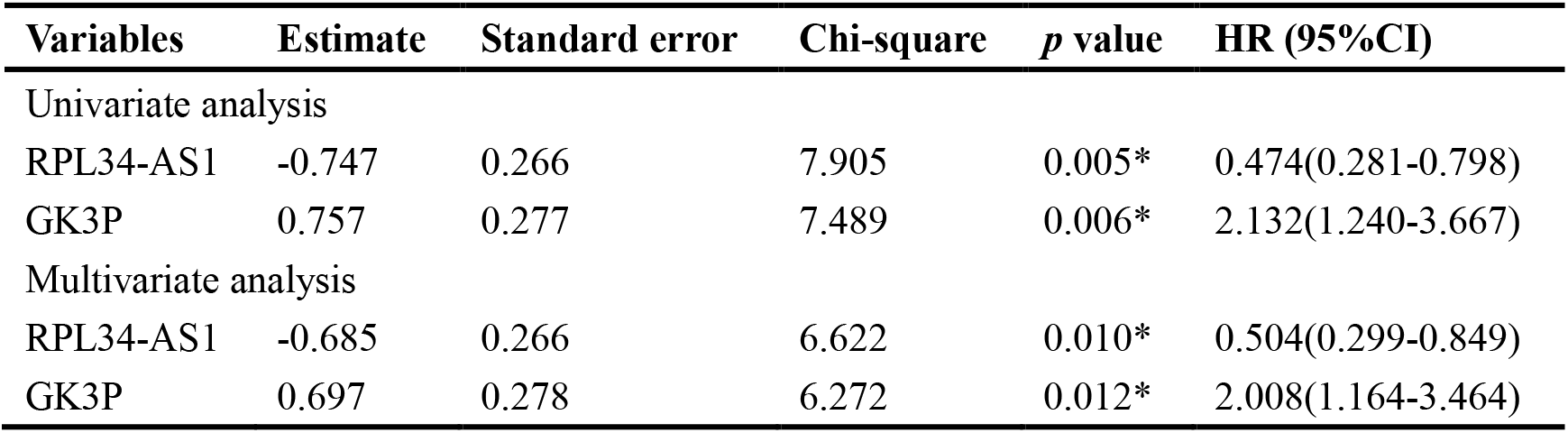
Prognostic value of the differentially expressed lncRNAs by univariate and multivariate Cox regression analyses.

A risk-score formula was established based on the combination of the expression profiles of each prognostic lncRNA, weighted by the regression coefficients that were calculated by multivariate Cox regression analysis. The formula was listed as follows: *Risk score* = *exp_RPL34_*_−*AS*1_ *(−0.685) + *exp_GK3p_* * (0.697). We calculated the risk score of the two-lncRNA expression signature for each patient.

Based on the median risk score as a cutoff value, all EC patients were divided into two groups: the low-risk score group (n = 75) and the high-risk score group (n = 75) (Fig 4). Meanwhile, K-M curves confirmed that the mean survival time (and standard deviation) of patients in the low-risk score group was 625.560 (± 507.079) days, significantly better than patients in the high-risk score group (478.000 ± 427.816 days, *p* = 0.002) (Fig 5A). Furthermore, this risk score model could be used to predict 5-year survival of EC patients as the area under ROC curve was 0.647 (Fig 5B).

**Figure 4.**
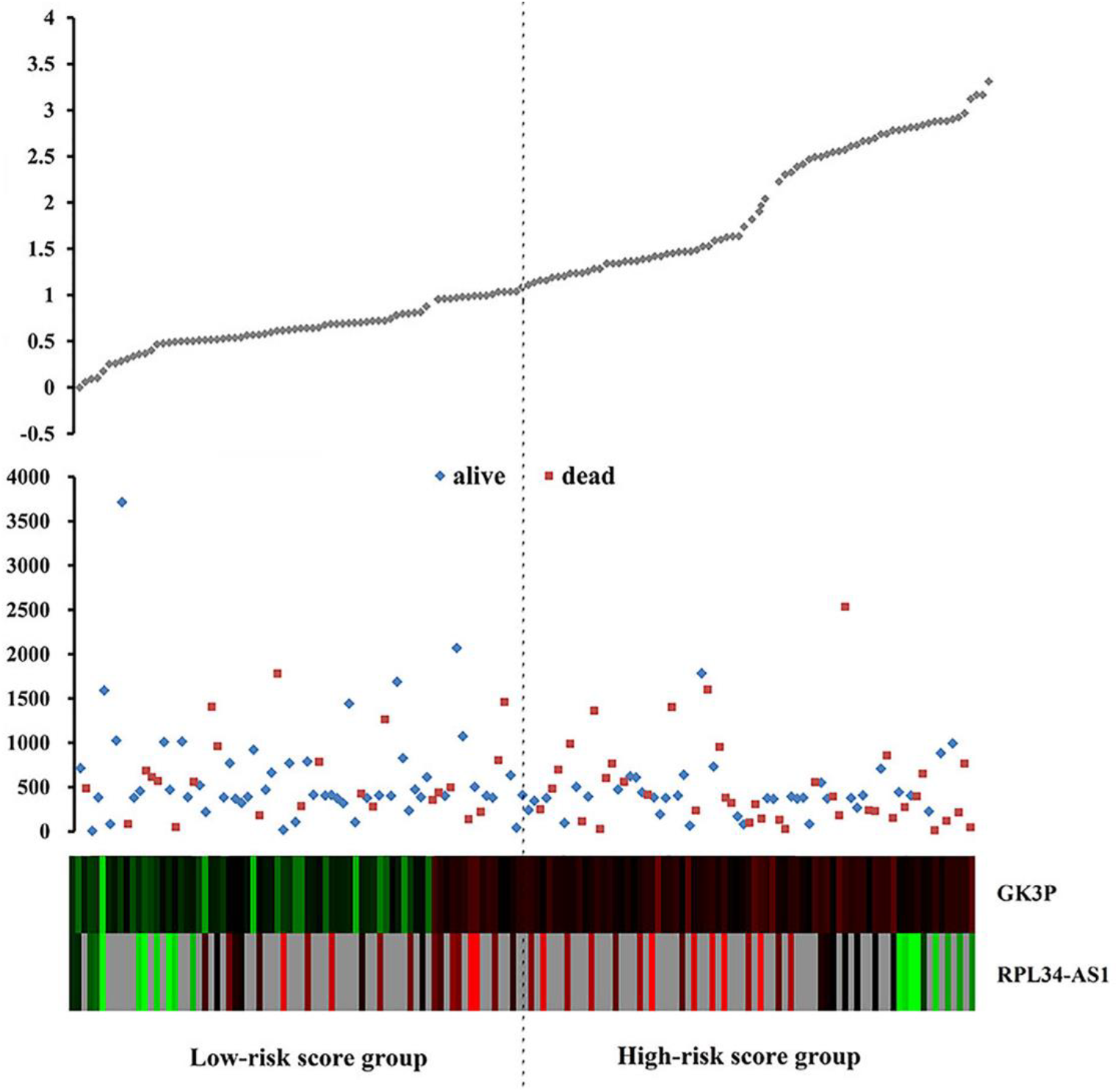
Expression of two lncRNAs, risk score distribution and survival in esophageal cancer patients. The risk scores for all patients are shown in the top; following up and survival of each patient are plotted in the middle; expression distribution of two lncRNAs by low-risk and high-risk score groups are shown in the bottom.

**Figure 5.**
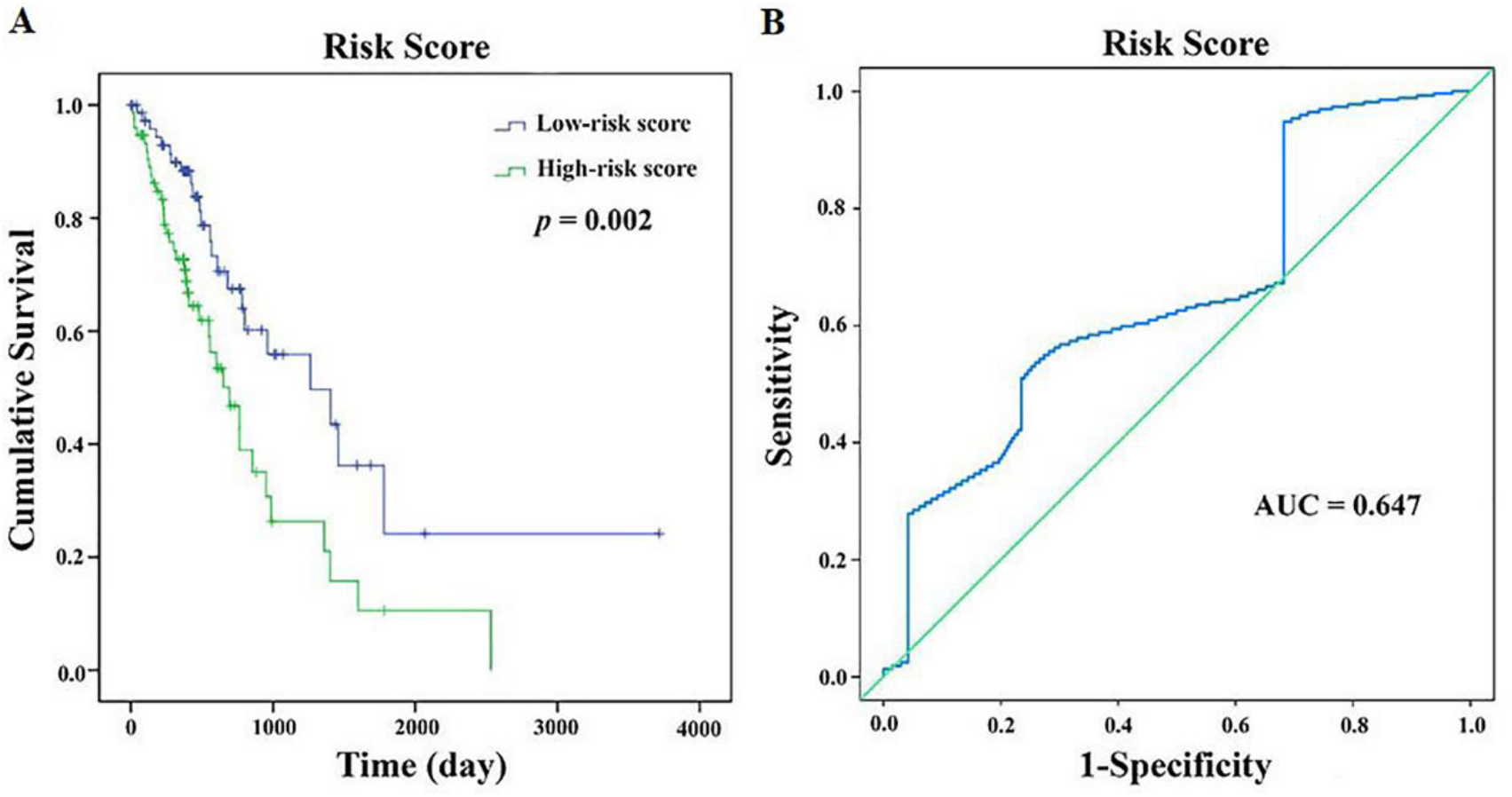
Risk score model for the outcome of esophageal cancer patients. (A) Kaplan-Meier curve tests the risk score model in overall survival; (B) Time-dependent ROC curve evaluates the efficacy of the risk score model in predicting 5-year survival.

The significant expression patterns of these two candidate lncRNAs in the tumor tissues/adjacent normal tissues group (Fig 6A), and in the low-risk score/high-risk score group were presented in Figure 6B.

**Figure 6.**
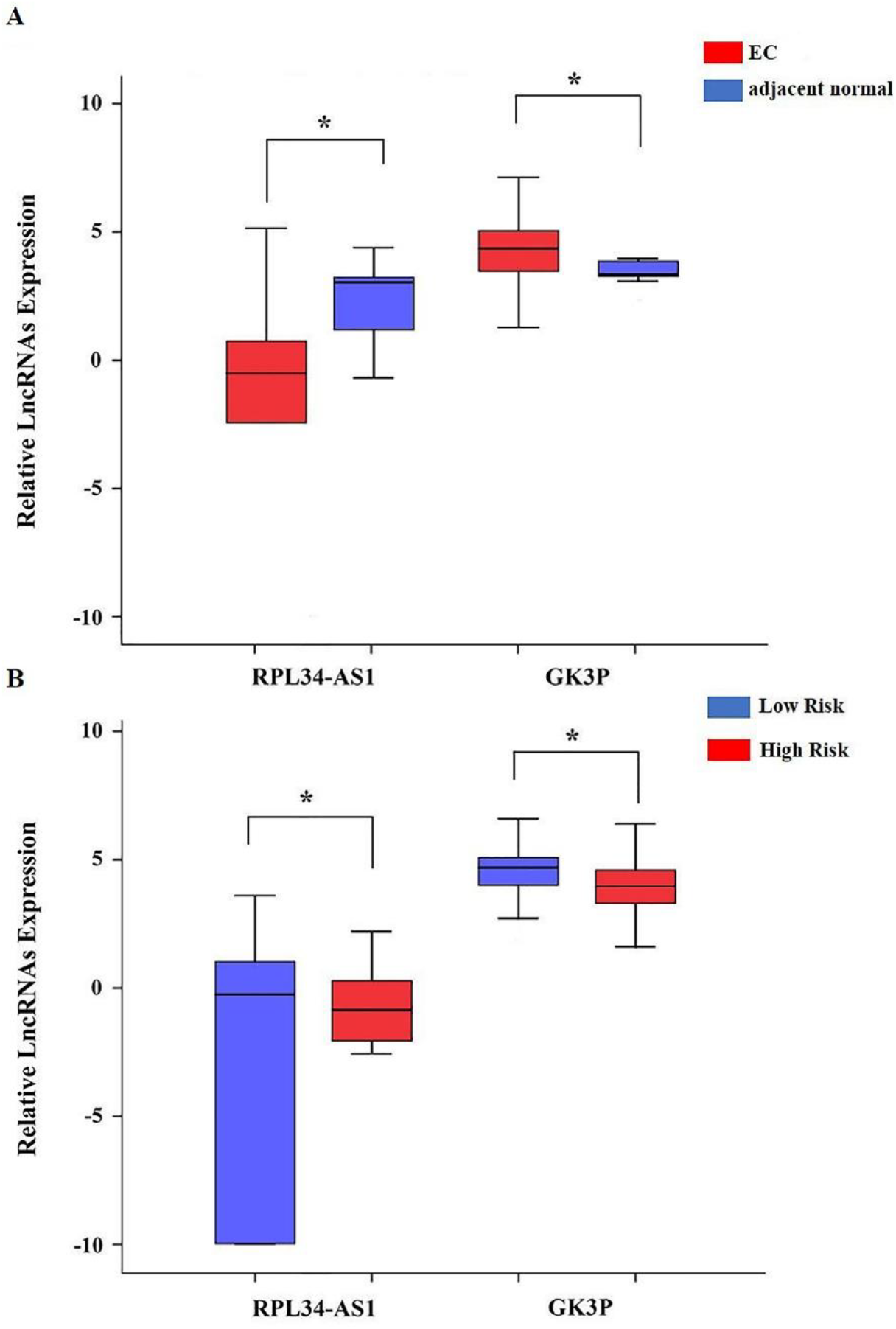
Comparison of the expression level of two lncRNAs. (A) The expression level of lncRNAs between tumor tissues and adjacent normal tissues; (B) The expression level of lncRNAs between low-risk and high-risk score groups.

### Correlation between the two-lncRNA signature and clinical characteristics

To verify the correlation between the two-lncRNA signature and clinical characteristics in EC patients, we used univariate Cox regression analysis and multivariate Cox regression analysis.

The univariate Cox regression analysis indicated that gender, TNM stage, N stage, M stage, additional treatment completion success outcome, residual tumor, and neoplasm tumor status could significantly predict poorer survival of EC patients (*p* < 0.05, Table 2). The multivariate Cox regression analysis showed that neoplasm tumor status (*p* = 0.036) and risk score (*p* = 0.001) could serve as independent prognostic factors in EC patients (Table 2).

**Table 2.** Predictive values of the related clinical characteristics and risk score.

| Variables |  | Univariate analysis |  | Multivariate analysis |  |
| --- | --- | --- | --- | --- | --- |
|  |  | HR (95% CI) | *p* value | HR (95% CI) | *p* value |
| Race | White | 1(reference) |  |  |  |
|  | Black | 0.931(0.126-6.869) | 0.944 |  |  |
|  | Asian | 0.744(0.347-1.597) | 0.448 |  |  |
| Gender | Female | 1(reference) |  |  |  |
|  | Male | 2.961(1.069-8.207) | 0.037* |  |  |
| Age | <=65 | 1(reference) |  |  |  |
|  | >65 | 1.326(0.791-2.223) | 0.284 |  |  |
| TNM stage | I | 1(reference) |  | 1(references) |  |
|  | II | 1.600(0.603-4.246) | 0.345 | 1.434(0.315-6.530) | 0.642 |
|  | III | 4.128(1.510-11.282) | 0.006* | 5.852(1.295-26.442) | 0.022* |
|  | IV | 8.087(2.441-26.794) | 0.001* | 3.055(0.473-19.733) | 0.241 |
| T stage | T1 | 1(reference) |  |  |  |
|  | T2 | 0.916(0.439-1.908) | 0.814 |  |  |
|  | T3 | 1.212(0.621-2.365) | 0.573 |  |  |
|  | T4 | 2.510(0.697-9.043) | 0.159 |  |  |
| N stage | N0 | 1(reference) |  |  |  |
|  | N1 | 2.272(1.274-4.051) | 0.005* | 1.718(0.589-5.008) | 0.258 |
|  | N2 | 3.748(1.371-10.246) | 0.010* | 0.866(0.173-4.342) | 0.230 |
|  | N3 | 3.042(1.019-9.086) | 0.046* | 2.287(0.351-14.914) | 0.820 |
| M stage | M0 | 1(reference) |  |  |  |
|  | M1 | 3.760(1.740-8.124) | 0.001* |  |  |
| Neoplasm histologic grade | G1 | 1(reference) |  | 1(references) |  |
|  | G2 | 2.341(0.800-6.852) | 0.121 | 1.262(0.300-5.309) | 0.453 |
|  | G3 | 2.150(0.720-6.425) | 0.170 | 0.903(0.212-3.849) | 0.481 |
| Additional treatment completion success outcome | CR | 1(reference) |  |  |  |
|  | SD | 1.079(0.137-8.490) | 0.943 |  |  |
|  | PD | 2.452(1.013-5.940) | 0.047* |  |  |
| Radiotherapy | No | 1(reference) |  |  |  |
|  | Yes | 0.834(0.411-1.689) | 0.614 |  |  |
| Columnar mucosa dysplasia | No | 1(reference) |  |  |  |
|  | Low | 3.558(0.952-13.300) | 0.059 |  |  |
|  | High | 1.229(0.554-2.728) | 0.613 |  |  |
| Lymph node metastasis | No | 1(reference) |  | 1(reference) |  |
|  | Yes | 1.674(0.951-2.945) | 0.074 | 0.663(0.279-1.575) | 0.274 |
| Residual tumor | R0 | 1(reference) |  | 1(reference) |  |
|  | R1+R2 | 2.523(1.290-4.933) | 0.007* | 2.152(0.829-5.581) | 0.107 |
| Neoplasm tumor status | Tumor free | 1(reference) |  | 1(reference) |  |
|  | With tumor | 2.483(1.451-4.248) | 0.001* | 2.132(1.331-5.766) | 0.036* |
| Risk score | Low | 1(reference) |  | 1(reference) |  |
|  | High | 2.210(1.312-3.720) | 0.003* | 3.500(1.685-7.274) | 0.001* |

The K-M curves of these clinical characteristics indicated that TNM stage (*p* < 0.001), N stage (*p* = 0.001), M stage (*p* < 0.001), and residual tumor (*p* = 0.005), and neoplasm tumor status (*p* = 0.001) were significantly associated with OS of EC patients (Fig 7A). Moreover, we assessed the association between the risk score and clinical characteristics and found that the risk score might have the prognostic significance in predicting some clinical characteristics, including TNM stage (AUC = 0.643, *p* = 0.040) and N stage (AUC = 0.579, *p* = 0.032) (Fig 7B).

**Figure 7.**
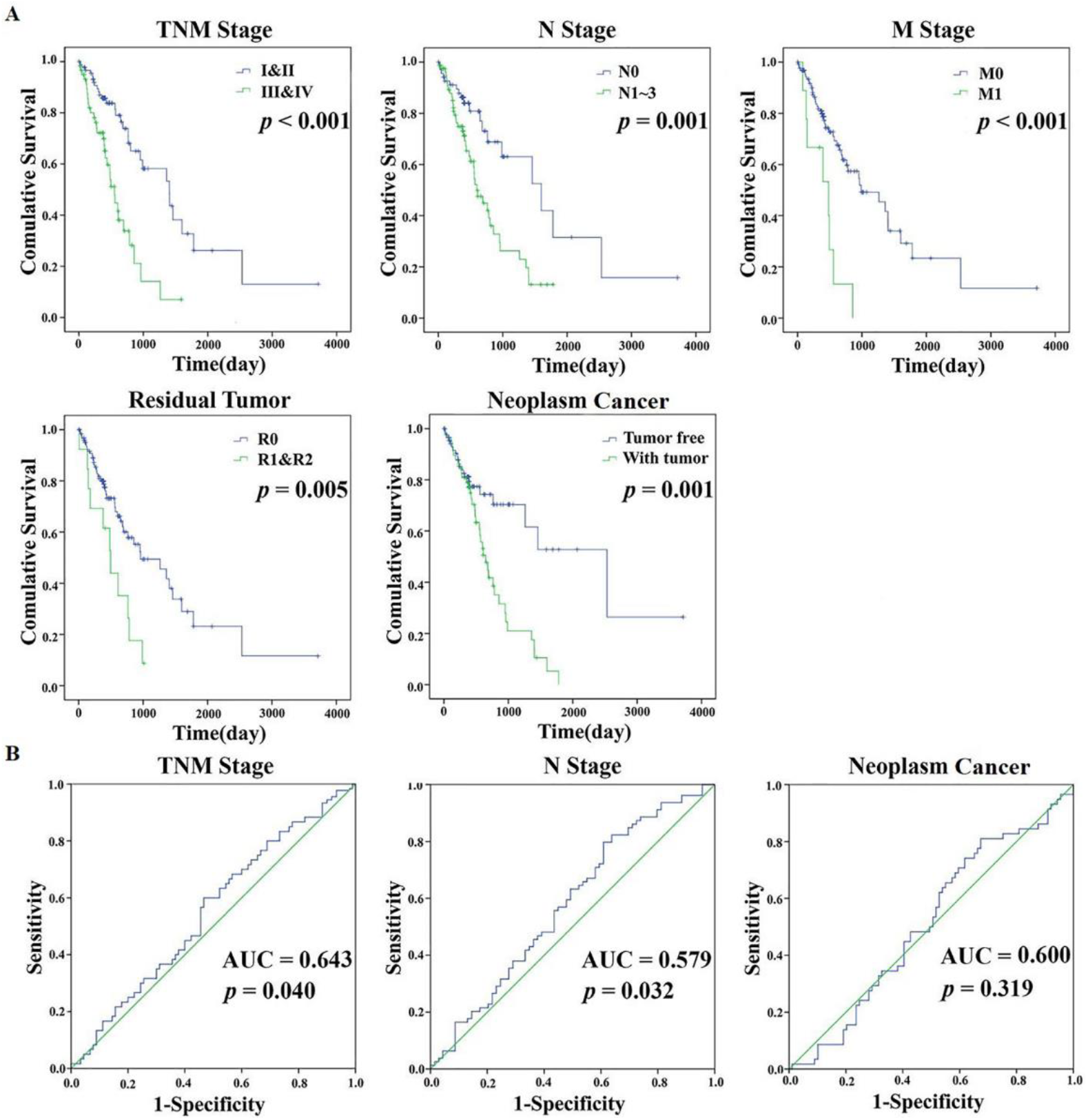
Prognostic value of the clinical characteristics in overall survival and predictive value of the risk score model for specific clinical characteristics. (A) Kaplan-Meier curves reveal five independent prognostic indicators; (B) ROC curves assess the risk score model in predicting clinical characteristics.

### Functional assessment of the two-lncRNA signature

629 genes (extracted from TCGA database) were co-expressed with these two lncRNAs (Pearson |R| > 0.5), including 622 genes co-expressed with RPL34-AS1 and seven genes co-expressed with GK3P (Supplemental Table 1). We identified 89 enriched GO terms and seven pathways (*p* <0.05, enrichment score > 1.5, Supplemental Table 2) by gene enrichment and functional annotation analysis. The top GO biological process and KEGG pathway (Table 3 and Fig 8) were detection of chemical stimulus involved in sensory perception of smell (GO: 0050911) and olfactory transduction (hsa: 04740), respectively. Finally, we constructed a PPI network of 283 genes that were considered as hub genes (Fig 9).

**Figure 8.**
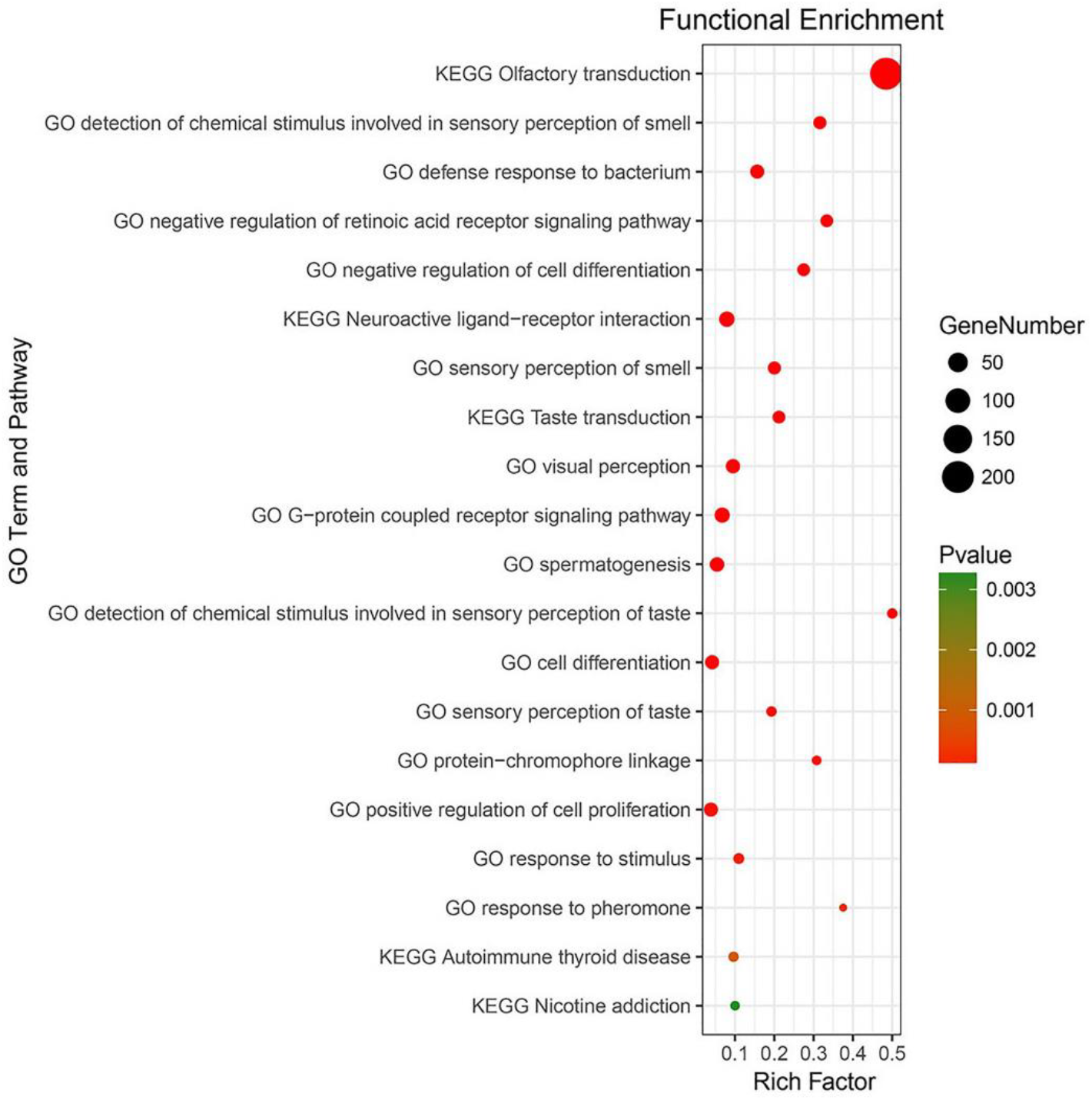
Top 20 enrichment of GO terms and KEGG pathways of co-expressed mRNAs.

**Figure 9.**
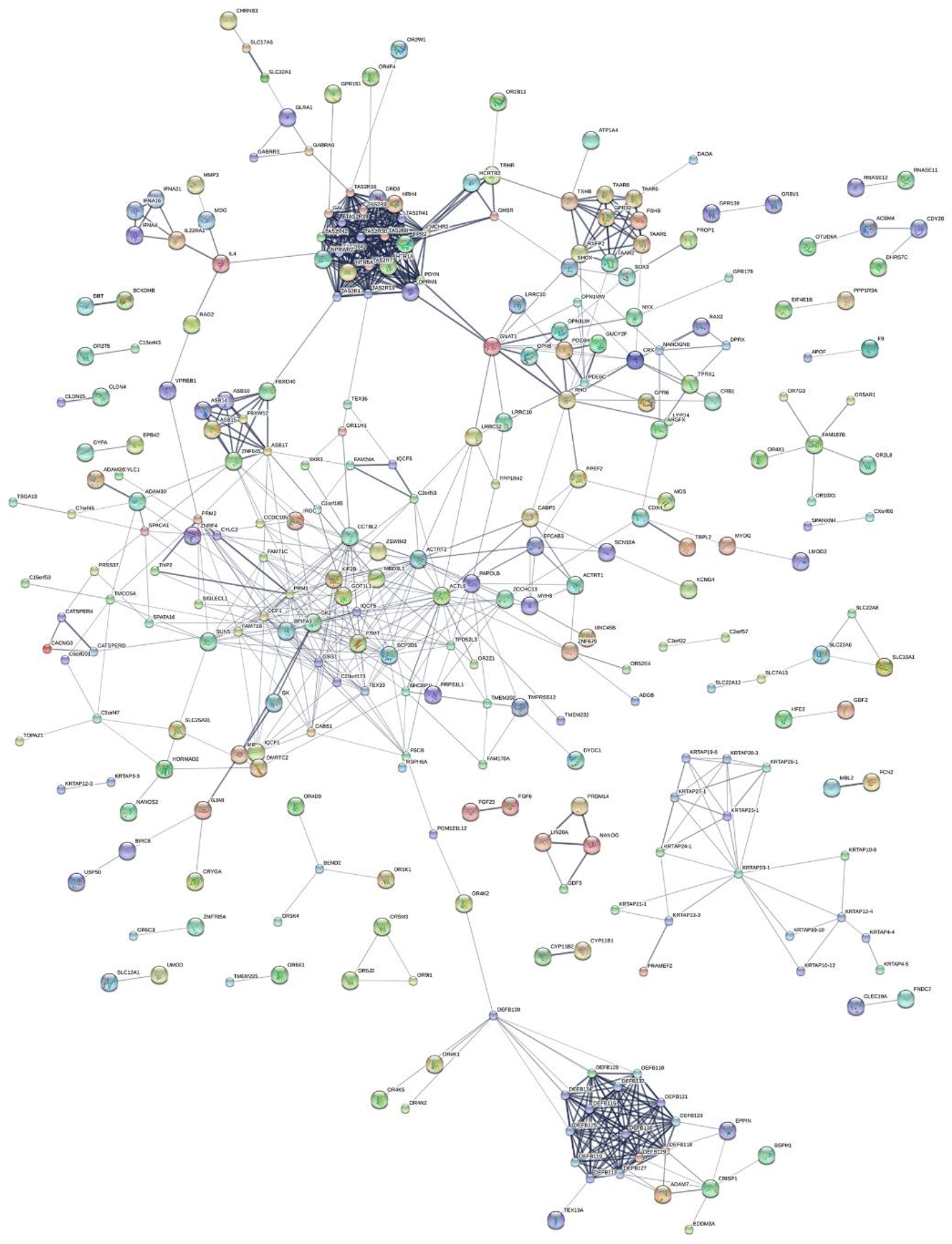
Map presented the PPI network of co-expressed genes.

**Table 3.** Top 20 KEGG and GO terms enriched by the coding genes.

| ID | Biological process and pathway | Count | Enrichment | -lg *P* | -lg FDR |
| --- | --- | --- | --- | --- | --- |
| **GO** |  |  |  |  |  |
| GO:0050911 | detection of chemical stimulus involved in sensory perception of smell | 12 | 30.048 | 13.860 | 11.040 |
| GO:0042742 | defense response to bacterium | 16 | 14.926 | 13.219 | 10.700 |
| GO:0048387 | negative regulation of retinoic acid receptor signaling pathway | 11 | 31.717 | 13.002 | 10.660 |
| GO:0045596 | negative regulation of cell differentiation | 11 | 26.167 | 11.954 | 9.737 |
| GO:0007608 | sensory perception of smell | 12 | 19.030 | 11.238 | 9.117 |
| GO:0007601 | visual perception | 17 | 9.037 | 10.400 | 8.359 |
| GO:0007186 | G-protein coupled receptor signaling pathway | 21 | 6.384 | 9.925 | 7.950 |
| GO:0007283 | spermatogenesis | 18 | 5.143 | 7.047 | 5.131 |
| GO:0050912 | detection of chemical stimulus involved in sensory perception of taste | 5 | 47.576 | 6.914 | 5.049 |
| GO:0030154 | cell differentiation | 16 | 3.985 | 4.815 | 2.995 |
| GO:0050909 | sensory perception of taste | 5 | 18.298 | 4.558 | 2.780 |
| GO:0018298 | protein-chromophore linkage | 4 | 29.277 | 4.494 | 2.754 |
| GO:0008284 | positive regulation of cell proliferation | 16 | 3.704 | 4.417 | 2.712 |
| **KEGG** |  |  |  |  |  |
| hsa: 04740 | Olfactory transduction | 201 | 46.085 | 289.793 | 287.644 |
| hsa: 04080 | Neuroactive ligand-receptor interaction | 22 | 7.557 | 11.850 | 10.002 |
| hsa: 04742 | Taste transduction | 11 | 20.128 | 10.586 | 8.914 |
| hsa: 05320 | Autoimmune thyroid disease | 5 | 9.149 | 3.059 | 1.512 |
| hsa: 05033 | Nicotine addiction | 4 | 9.515 | 2.485 | 1.035 |
| hsa: 04744 | Phototransduction | 3 | 10.195 | 1.905 | 0.534 |
| hsa: 04140 | Regulation of autophagy | 3 | 7.319 | 1.496 | 0.192 |

## DISCUSSION

Esophageal cancer (EC) is among the deadliest malignancies in the world, with high morbidity and mortality (Griffin & Berrisford 2013; Torre et al. 2015). The onset of dysphagia is associated symptom of EC, and a small proportion of patients with EC survive for five years (di Pietro et al. 2017). The novel biomarker for early diagnosis, process monitoring, and prognostic evaluation might increase the survival rate for EC patients. In the past decade, researchers have reported that about 70% of the genome is transcribed in various cell types and contexts (Djebali et al. 2012; Okazaki et al. 2002), nearly 80% of the genome is biochemically active (2012), and many DNAs code for RNA as end products instead of proteins (Pennisi 2012). LncRNAs are engaged in a wide range of cellular mechanisms, such as DNA methylation and histone modifications (O’Leary et al. 2015; Plath et al. 2003). In addition, increasing evidence has proved that lncRNAs function as a critical player in gene regulation and trend to be expressed in a more tissue- and cell-specific manner, which demonstrated the crucial advantages of lncRNAs as diagnostic or prognostic biomarkers in cancers (Bolha et al. 2017; Evans et al. 2016; Yang et al. 2014). Therefore, development of novel lncRNA-related biomarkers in prognostic prediction of EC patients may improve treatment decisions regarding the aggressiveness or recurrence of the disease. In the present study, we investigated the impact of lncRNAs on EC patients’ survival, which were based on RNA sequencing technology.

EC usually presents late and carries a poor prognosis (Bird-Lieberman & Fitzgerald 2009; Meves et al. 2015). Therefore, we aimed to find a novel lncRNA signature and to investigate whether it affects the prognosis of all stages of EC patients. In this study, we first screened out the differentially expressed lncRNAs in tumor tissues, by comparing with the expression in non-tumor tissues. Among candidate lncRNAs, we identified two lncRNAs as having significant prognostic value for survival in all EC patients by using univariate and multivariate Cox regression models. Then we constructed a risk-score model by combining the expressions of these two lncRNAs and found that the two-lncRNA expression signature could predict OS for all EC patients.

Recently, Fan and Liu identified an eight-lncRNA signature (GS1-600G8.5, CTD-2357A8.3, LINC00365, RP11-705O24.1, RP1-90J4.1, RP11-327J17.1, LINC01554,and LINC00176) which might be able to predict survival of EC patients (Fan & Liu 2016). Compared to their work, our study used “Fold change > 2 or < 0.5, *p* < 0.05, and FDR < 0.05” as the screening criteria, which was more rigorous for bioinformatics analysis. Therefore, the number of candidate prognostic lncRNAs in both studies were different. Moreover, we combined RNA sequencing data with clinical information and found that this two-lncRNA signature could be served as an indicator of specific clinical characteristics (TNM stage and N stage) for EC patients.

A series of evidence-based papers have proved that lncRNA could serve as a carcinogen or tumor suppressor of EC. Despite this, the role of most lncRNAs in EC remains unknown. Lin et al. reported that HOXA13 might promote metastasis and tumorigenesis in EC cells (Lin et al. 2017). Yao et al. found that RP11-766N7.4 was associated with carcinogenesis of EC (Yao et al. 2017). Furthermore, Wu et al. suggested that NORAD was related to poor prognosis in EC patients (Wu et al. 2017).

Regarding these two lncRNAs, Zhao et al. reported that the decreased expression of RPL34-AS1 was correlated with larger gastric tumors (Zhao et al. 2015). For GK3P, it has not been reported yet. Besides, the biological functions of these two lncRNAs have not been investigated in EC. Here, we assessed the functional relevance of these two lncRNAs by using DAVID bioinformatics resources. 629 genes were identified to have significant co-expression with these two lncRNAs. The relevant genes were mainly enriched in olfactory transduction, neuroactive ligand-receptor interaction, detection of chemical stimulus involved in sensory perception of smell, and defense response to bacterium. Additionally, 283 genes were identified as hub genes regulated by these two lncRNAs via the PPI network.

The findings of our study may have essential clinical significance. However, some limitations should be taken into consideration as well. Firstly, the patient cohort of TCGA was obtained from multi-institutional sites, and the results still need to be validated in other cohorts in the future study. Secondly, only lncRNAs were involved in our work, and this study could not represent the whole transcription alteration associated with EC. Thirdly, the functional role of RPL34-AS1 and GK3P are still unknown. Thus, we need well-designed experiments to validate the biological functions of these two lncRNAs in EC.

## CONCLUSIONS

We identified two lncRNAs associated with the survival of EC patients in a large cohort from the TCGA database and developed a lncRNA signature. Further analysis indicated that the two-lncRNA signature could be a prognostic indicator independent of clinical characteristics. It can serve as a novel biomarker of prognostic prediction for EC patients. Further functional validations are needed to explore the regulatory mechanisms of these lncRNAs in EC.

## ACKNOWLEDGMENTS

We thank Mr. Dong-Ling Chen for his help in technical assistance.

